# Entropy-Based Analysis of Vertebrate Sperm Protamine Sequences: Evidence of Dityrosine and Cysteine-Tyrosine Cross-Linking in Sperm Protamines

**DOI:** 10.1101/845180

**Authors:** Christian D. Powell, Daniel C. Kirchhoff, Jason E. DeRouchey, Hunter N.B. Moseley

## Abstract

**Background:** Spermatogenesis is the process by which germ cells develop into spermatozoa in the testis. Sperm protamines are small, arginine-rich nuclear proteins which replace somatic histones during spermatogenesis, allowing a hypercondensed DNA state that leads to a smaller nucleus and facilitating sperm head formation. In eutherian mammals, the protamine-DNA complex is achieved through a combination of intra- and intermolecular cysteine cross-linking and possibly histidine-cysteine zinc ion binding. Most metatherian sperm protamines lack cysteine but perform the same function. This lack of dicysteine cross-linking has made the mechanism behind metatherian protamines folding unclear.

**Results:** Protamine sequences from UniProt’s databases were pulled down and sorted into homologous groups. Multiple sequence alignments were then generated and a gap weighted relative entropy score calculated for each position. For the eutherian alignments, the cysteine containing positions were the most highly conserved. For the metatherian alignment, the tyrosine containing positions were the most highly conserved and corresponded to the cysteine positions in the eutherian alignment.

**Conclusions:** High conservation indicates likely functionally/structurally important residues at these positions in the metatherian protamines and the correspondence with cysteine positions within the eutherian alignment implies a similarity in function. One explanation is that the metatherian protamine structure relies upon dityrosine cross-linking between these highly conserved tyrosines. Also, the human protamine P1 sequence has a tyrosine substitution in a position expecting eutherian dicysteine cross-linking. Similarly, some members of the metatherian Planigales genus contain cysteine substitutions in positions expecting plausible metatherian dityrosine cross-linking. Rare cysteine-tyrosine cross-linking could explain both observations.

## Background

The process in which male germ cells develop into sperm cells is called spermatogenesis. During spermatogenesis, DNA undergoes hypercondensation in order to form a smaller nucleus. This is accomplished through the replacement of a vast majority of somatic DNA histones (>90%) with one of three nuclear proteins; sperm-specific histones, protamine-like proteins, or protamines [Balhorn 2007]. In mammals, sperm protamines are small (<60 amino acids), arginine-rich nuclear proteins. Hypercondensation of DNA mediated by protamines result in haploid male germ cell nuclei, which are genetically inactive but just 1/20th the size of a somatic cell nucleus [Steger & Balhorn 2018]. This reorganization of the spermatozoa DNA is also thought to protect the paternal genome against oxidative damage [Balhorn 2007; Bennetts & Aitken 2005; Villani et al. 2010; Enciso et al. 2011].

Genetically the family of sperm protamines is highly diverse, being observed across the tree of life. For example, a single species of fish can contain multiple genes for protamine and protamine-like proteins, whereas birds tend to have two identical copies of a single protamine gene [Balhorn 2007]. While all protamines perform the task of binding and condensing DNA, the sizes and structural components of the protamines can vary greatly from species to species. Despite these differences most all sperm protamines include large arginine-rich DNA binding regions and phosphorylation sites [Balhorn 2007; Biegeleisen 2006; Queralt et al. 1995]. The positively charged arginine residues in these binding regions are able to engage in an electrostatic interaction with the negatively charged DNA phosphate backbone [Biegeleisen 2006]. These interactions form a toroid shaped protamine-DNA complex conforming to an internal hexagonal lattice [Brewer 2011]. The various phosphorylation sites in the protamine sequences are involved in a number of post-translational modifications and are thought to regulate the interactions with DNA.

Some of the simplest protamines are those of fish. Like most sperm protamines, the sequences of fish protamines are populated with a large percentage of arginine residues. However, fish protamines tend to be under 35 amino acids in length and contain increased frequencies of arginine (approximately 70%) in comparison to their mammalian analogs. The secondary structure of the protamines consist of multiple beta turns, with limited CD, NMR, and fluorescence data, indicating the formation of a possible globular structure [Arellano et al. 1988; Cid & Arellano 1982].

Mammals have a relatively conserved set of protamines, with metatherian mammals having only one protamine gene, while eutherian mammals have two to three varieties. These mammalian sequences tend to start with MARYR at the N-terminus, typically followed by a region containing a phosphorylation site (or multiple phosphorylation sites in eutherian sperm protamine P2s), then a DNA binding region comprised of multiple blocks of arginine residues, and ending with a varied C-terminal region [Queralt et al. 1995].

The eutherian protamine P1 is encoded by the PRM1 gene. Alignment of the sequences of eutherian mammal sperm protamine P1 have shown the sequences to be relatively conserved. In eutherian sperm protamine P1 sequences, the arginine-rich DNA binding regions are broken up by cysteine residues, which are involved in both inter- and intramolecular disulfide cross-linkings [Balhorn et al. 1991; Queralt et al. 1995; Vilfan et al. 2004]. In bull protamine P1, the intra-protamine disulfide bonds were shown to create a hairpin-like structure, with disulfide crosslinks formed between the cysteines in positions 7 and 15 as well as the cysteines at positions 40 and 48. The remaining cysteine positions in bull protamine P1 are involved in inter-protamine bonding. More recently, we have shown that this disulfide mediated secondary structure of the bull protamine is required for proper chromatin remodeling [Hutchison et al. 2017; Kirchhoff et al. 2019].

The other two eutherian sperm protamine types are encoded in the PRM2 gene. These other protamine proteins are longer than the eutherian sperm protamine P1 type protamine and include a number of post-translational truncation sites in the N-terminal tail [Balhorn 2007]. Unlike the eutherian P1 type sperm protamines, the P2 protamines engage in zinc ion binding that is stochiometrically 1:1 for many eutherian mammals [Bench et al. 2000]. This zinc ion binding is achieved with highly conserved cysteine and histidine residues in the P2 protamine sequence. Both eutherian P1 and P2 type protamines engage in intermolecular disulfide cross-linking with one another when forming the DNA protamine complex [Vilfan et al. 2004]. For all eutherian sperm protamine types, it is thought that the cysteine cross-linkages are important for protecting the spermatozoa from oxidative damage [Bennetts & Aitken 2005; Villani et al. 2010; Enciso et al. 2011].

Also in eutherian mammals, a testis-specific variant of glutathione peroxidase (GPx4) is involved in the formation of the thiol cross-linking between and within the protamines and with protecting the sperm cells from oxidative stress due to reactive oxygen species [Pfeifer et al. 2001; Conrad et al. 2005]. In particular, Conrad et al. showed that without GPx4, sperm develop abnormal heads likely due to a lack of stabilizing disulfide cross-linking [Conrad et al. 2005].

In contrast to eutherian sperm protamines, little is known about metatherian sperm protamines, except that metatherian sperm protamines tend to lack cysteine residues with the only exception to this tendency involving species of the *Planigale* genus [Retief et al. 1995]. To our knowledge, there is no consensus on the structure of metatherian sperm protamines, nor is there prior evidence to suggest that inter- and intramolecular cross-linking occurs in metatherian sperm protamines, with the exception of species of the *Planigale* genus where it was suggested [Retief et al. 1995]. Additionally, it is unclear if GPx4 is required for the proper function of metatherian sperm protamines, although it is known that metatherian mammals do express glutathione peroxidase for defense against oxidative stress [Whittington et al. 1995]. Sequence data also exists for the testes specific version of glutathione peroxidase in Tasmanian Devils (G3WAH0_SARHA) [UniProt 2018]. Metatherian spermatozoa are more susceptible to oxidative damage, likely due to a lack of stabilizing disulfide cross-linkages [Bennetts & Aitken 2005; Villani et al. 2010; Enciso et al. 2011]. These current gaps in knowledge prompted the following analyses of multiple sequence alignments (MSAs) of eutherian P1, eutherian P2, metatherian P1, and fish sperm protamine sequences, which provide some insight into the structures of protamines and mechanism behind protamine mediated DNA condensation.

## Results

MSAs were generated for 145 eutherian sperm protamine P1, 16 eutherian sperm protamine P2, 95 metatherian sperm protamine, and 34 fish protamine sequences, all retrieved from the UniProt knowledgebase and aligned using MUSCLE 3.8.31 [UniProt 2018; Edgar 2004]. The MSAs were then analyzed using the relative entropy method described.

### Fish Protamine

From the fish protamine MSA, the relative entropy-based analysis showed that no position in the alignment had conservation scores above the relative entropy threshold. The most highly conserved positions were those containing only arginine residues. There was in fact a four-way tie for the most highly conserved position with positions 15, 16, 17, and 27 all having a conservation score equal to the conservation threshold of 4.135.

### Eutherian Sperm Protamine P1

For the eutherian sperm protamine P1 MSA, a total of nine positions were determined to be highly conserved, based on relative entropy scores and conservation threshold described above. All nine highly conserved positions within the alignments were positions comprised primarily of cysteine residues. In descending conservation score, the most highly conserved positions were positions 7, 49, 50, 60, 38, 17, 29, 6, and 37. All highly conserved positions within the alignment were composed of over 69% cysteine residues (excluding gaps) and the most highly conserved positions tended to all be within one residue of a known intramolecular cross-linking region (positions 7, 49, 50, 60, 17, and 6) [Balhorn et al. 1991].

### Eutherian Sperm Protamine P2

For the truncated eutherian sperm protamine P2 MSA, a total of eleven positions were determined to be highly conserved, based on relative entropy scores and conservation threshold described above. While intramolecular cross-linking in sperm protamine P2 proteins have yet to be determined [Vilfan et al. 2004], half of the positions identified as highly conserved in the P2 alignment consisted primarily of cysteine residues (positions 59, 75, 83, 93, and 107). The remaining positions are either primarily composed of histidine (positions 68, 89, 53, 85, and 110) or tyrosine (position 54) residues.

### Metatherian Sperm Protamine

When the relative entropy method was applied to the metatherian sperm protamine alignment, similar results are found in the eutherian sperm protamine P1 alignment. Instead of nine positions determined to be conserved, like in the eutherian P1 alignment, the metatherian alignment only has seven highly conserved positions. Of these seven positions, six were found to primarily contain tyrosine (positions 4, 57, 16, 62, 75, and 34). The remaining highly conserved position in the alignment primarily consisted of histidine residues (position 7).

### Whole Sequence Arginine-Lysine Density Analysis

The arginine-lysine frequencies for each protamine in each homologous protamine group are shown in Figure 5 and in Table 1. It is clear from the differences in these arginine-lysine frequency distributions that the relative proportion of the DNA binding region to the whole protamine sequence is quite different for the protamine groups, especially the eutherian P2 protamines.

**Table 1.** Arginine-Lysine Density of Sperm Protamine Groups. Low, quartile, and max information for eutherian P1, eutherian P2, truncated eutherian P2, metatherian, and fish protamine groups.

| Alignment | Min | Q1 | Median | Q3 | Max |
| --- | --- | --- | --- | --- | --- |
| Eutherian P1 | 0.429 | 0.471 | 0.520 | 0.580 | 0.652 |
| Eutherian P2 | 0.294 | 0.328 | 0.341 | 0.362 | 0.385 |
| Eutherian P2 Truncated | 0.473 | 0.509 | 0.545 | 0.559 | 0.589 |
| Metatherian | 0.478 | 0.587 | 0.600 | 0.624 | 0.656 |
| Fish | 0.562 | 0.636 | 0.652 | 0.667 | 0.704 |

**Figure 1.**
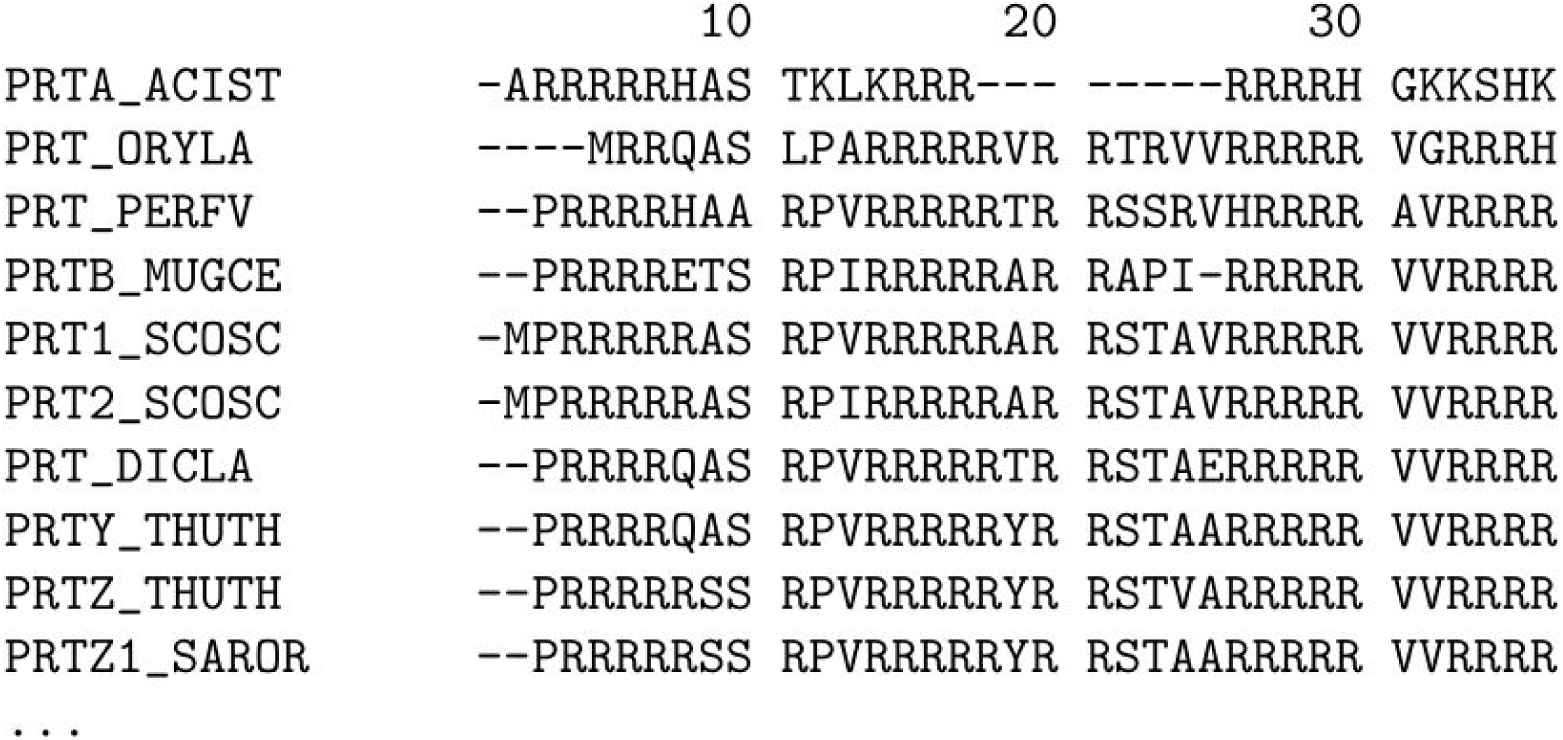
Alignment showing a selection of 10 fish protamine amino acid sequences from the alignment of fish protamine sequences. The full multiple sequence alignment used 34 sequences from the 2019_05 release of the UniProt knowledgebase and was aligned with MUSCLE 3.8.31. For full alignment, see Supplemental Figure 1.

**Figure 2.**
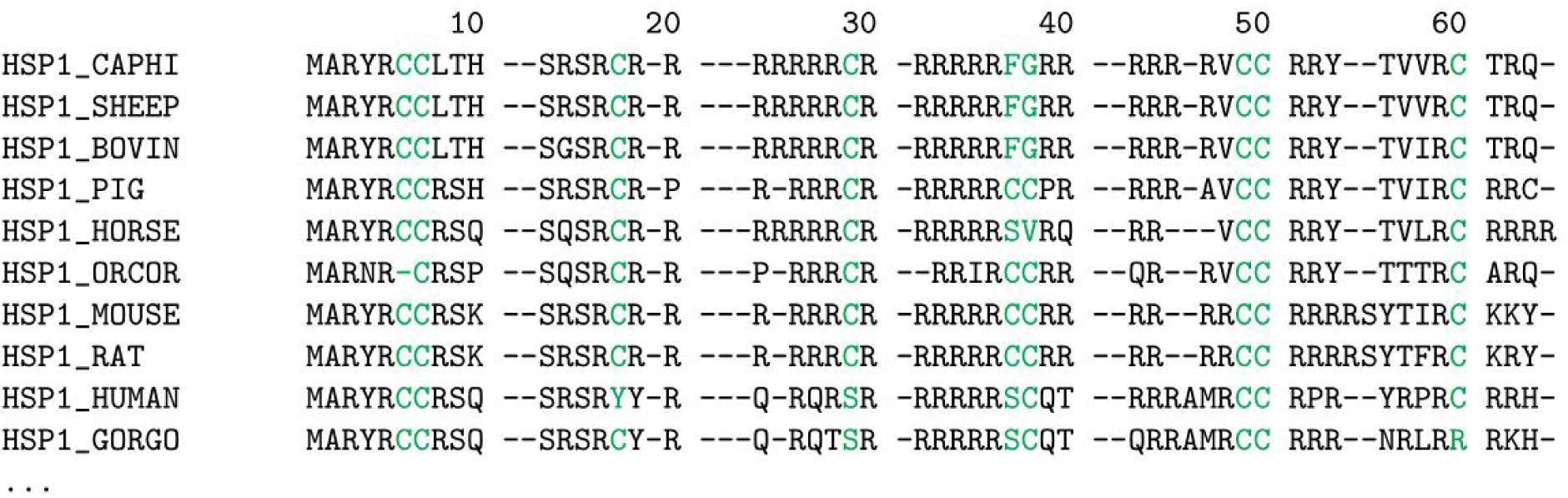
Alignment showing a selection of sperm protamine P1 amino acid sequences from 10 common eutherian mammals. The full multiple sequence alignment used 145 sequences from the 2019_05 release of the UniProt knowledgebase and was aligned with MUSCLE 3.8.31. Positions with relative entropy score greater than the conservation threshold of 4.135 are highlighted (see Supplemental Table 1). For full alignment see Supplemental Figure 2.

**Figure 3.**
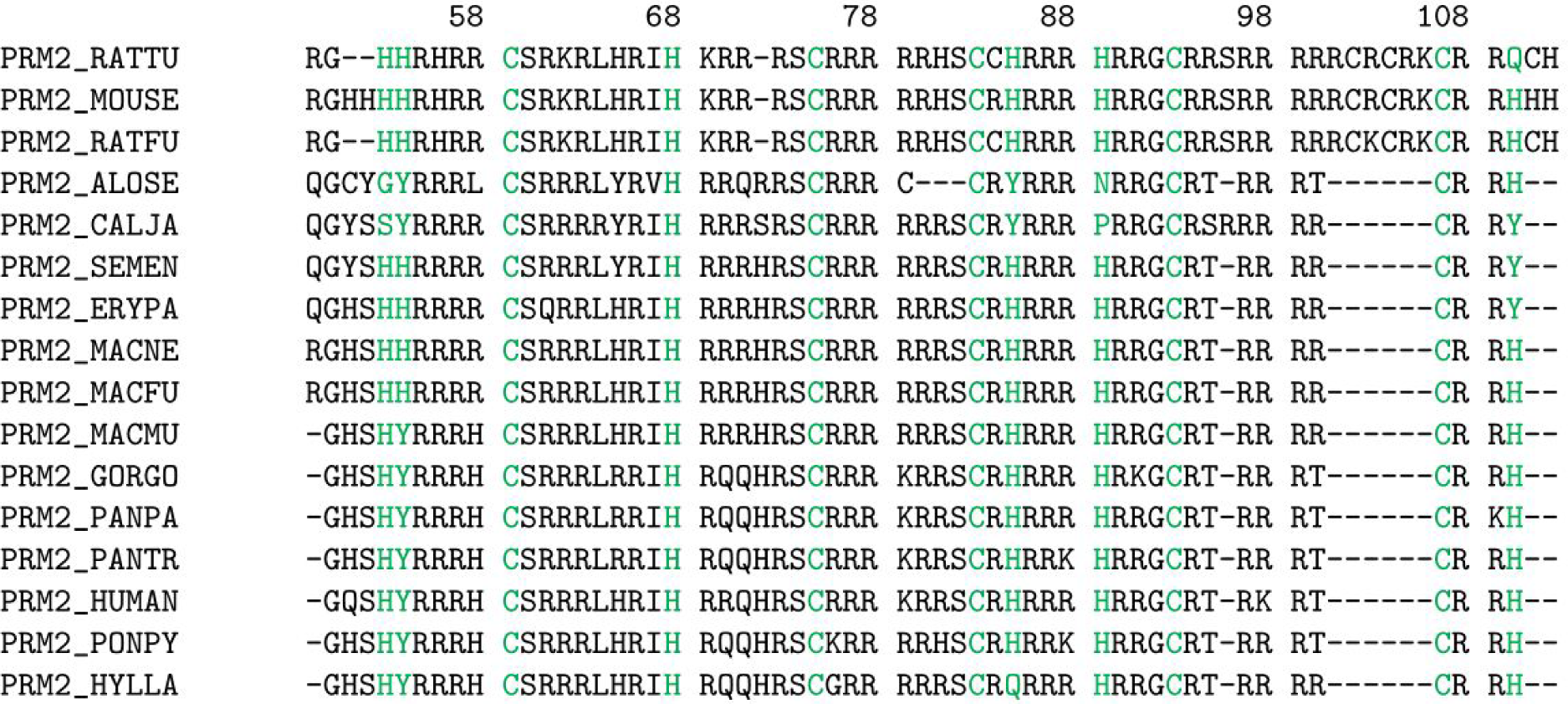
Alignment showing truncated sperm protamine P2 amino acid sequences from the alignment of 19 eutherian mammals. Sequences were pulled from the 2019_05 release of the UniProt knowledgebase and was aligned with MUSCLE 3.8.31. Positions with relative entropy score greater than the conservation threshold of 4.135 are highlighted (see Supplemental Table 2 for the truncated alignment and see Supplemental Table 3 for the untruncated alignment). For the untruncated alignment see Supplemental Figure 3.

**Figure 4.**
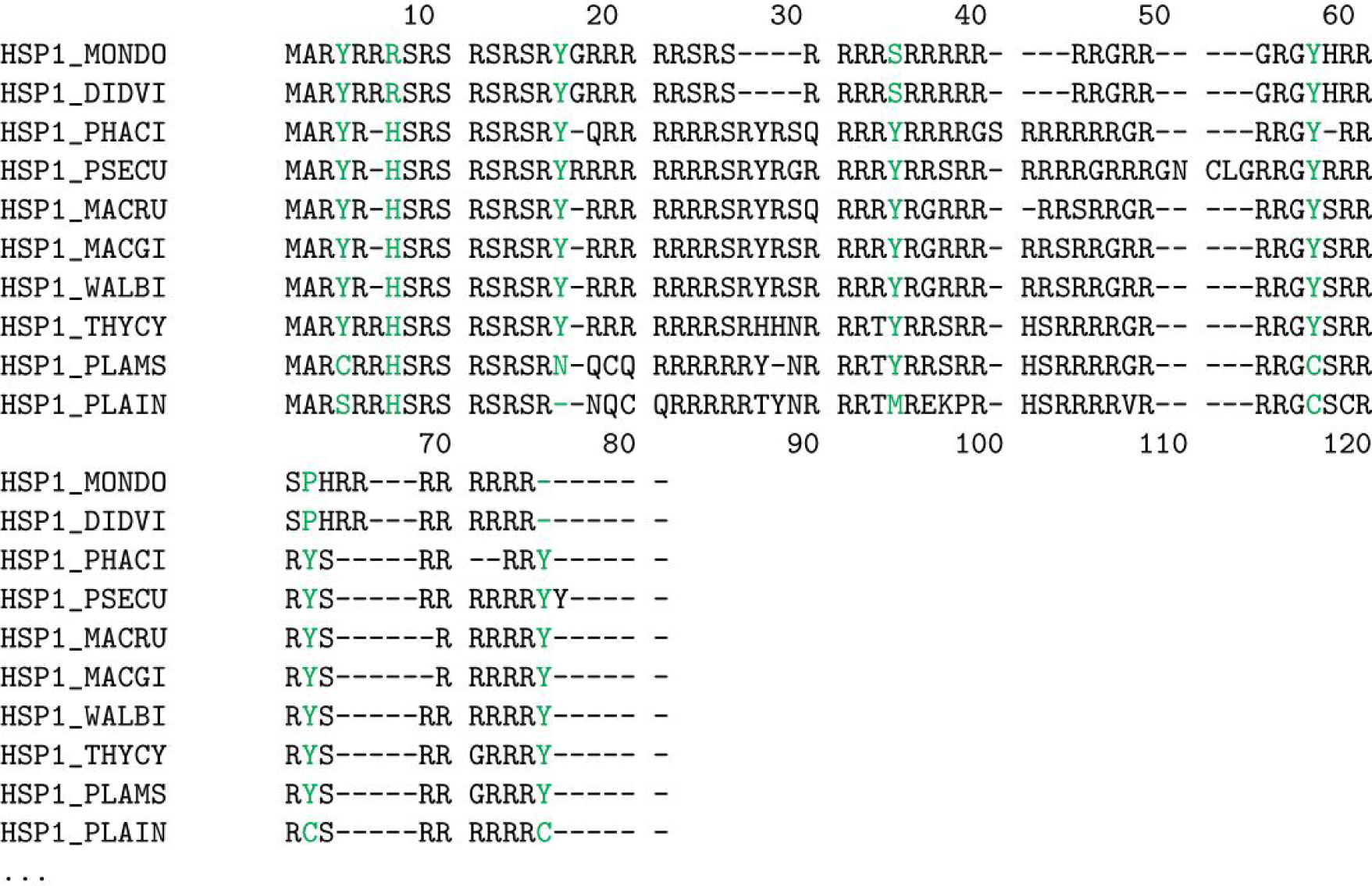
Alignment showing a selection of sperm protamine P1 amino acid sequences from 10 metatherian mammals. The full multiple sequence alignment used 95 sequences from the 2019_05 release of the UniProt knowledgebase and was aligned with MUSCLE 3.8.31. Positions with relative entropy score greater than the conservation threshold of 4.135 are highlighted (see Supplemental Table 5). For full alignment see Supplemental Figure 4.

**Figure 5.**
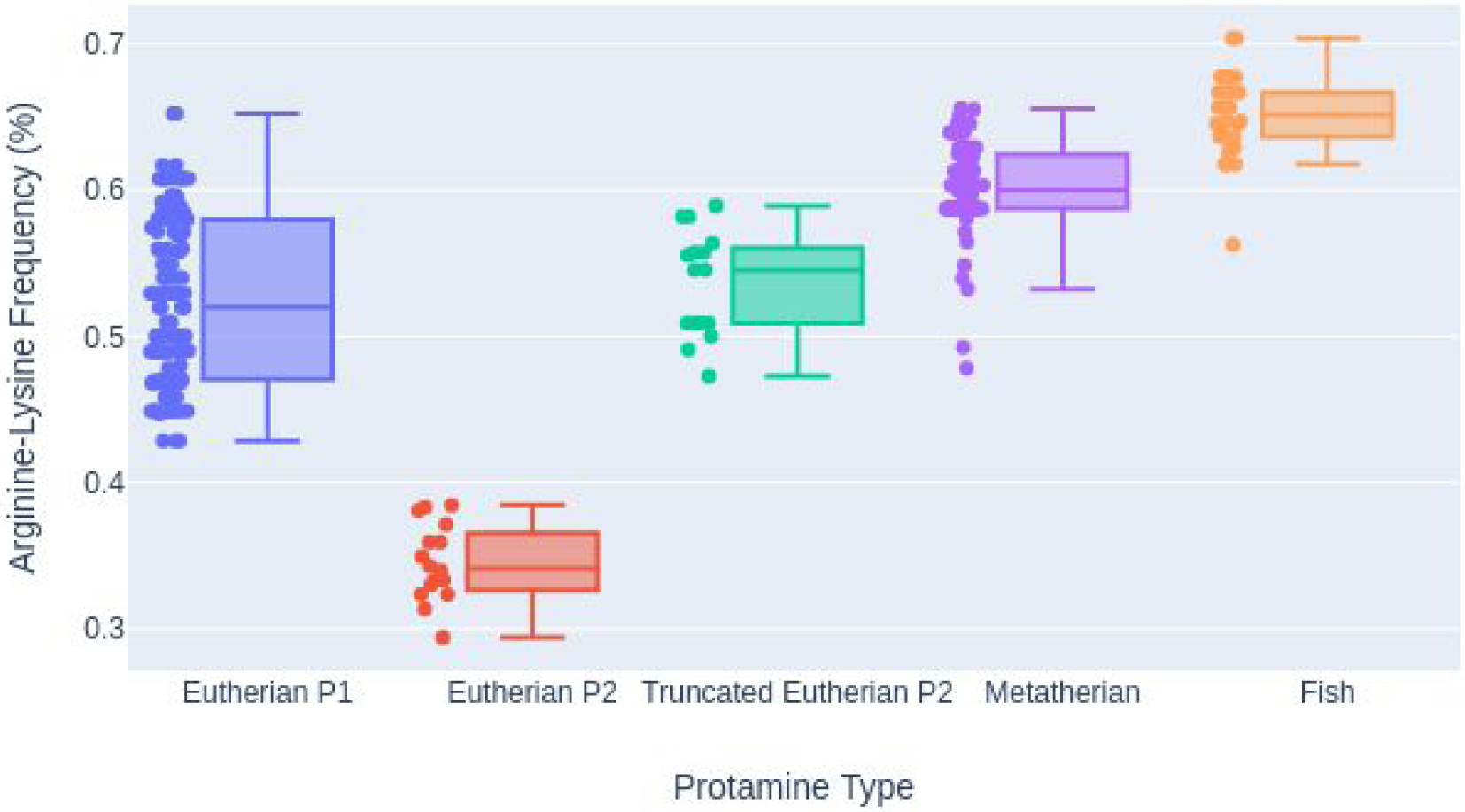
Box plots of the arginine frequency for eutherian protamine P1, truncated eutherian P2, metatherian P1, and fish protamines. Individual arginine frequencies are plotted next to the associated box plot.

### DNA Binding Region Arginine-Lysine Density Analysis

Figure 6 and Table 2 show the arginine-lysine frequencies in the hypothesized DNA binding regions for each protamine in the eutherian P1 and metatherian MSAs. FIsh protamines were included in Figure 6 for comparison. The distributions were analyzed using a Welch’s *t*-test which showed that each protamine group’s arginine-lysine frequency distribution in their hypothesized DNA binding region is statistically discrete from any other groups. Comparing the DNA binding region of the eutherian P1 sperm protamine group to that of the metatherian sperm protamine group yielded a p-value of 2.184e-2. Comparing the DNA binding region of the eutherian P1 sperm protamine group to the whole fish sequence sperm protamine group yielded a p-value of 4.762e-11. Comparing the DNA binding region of the metatherian sperm protamine group to the whole fish sequence sperm protamine group yielded a p-value of 4.987e-8. However, the differences between the medians of these distributions is less than 0.017 (see Table 2) and the possible functional consequences of these relatively small differences in arginine-lysine frequencies are unclear given the relatively high variance of each group. It is possible that the arginine-lysine frequency of the DNA binding region across all of the protamines is just a species- and protein-specific optimization of the DNA binding function.

**Table 2.** Arginine-Lysine Density of DNA Binding Regions. Low, quartile, and max information for the hypothesized DNA binding region of eutherian P1 and metatherian sperm protamines.

| Alignment | Min | Q1 | Median | Q3 | Max |
| --- | --- | --- | --- | --- | --- |
| Eutherian P1 DNA | 0.478 | 0.636 | 0.714 | 0.773 | 0.857 |
| Metatherian DNA | 0.529 | 0.667 | 0.697 | 0.719 | 0.781 |

**Figure 6.**
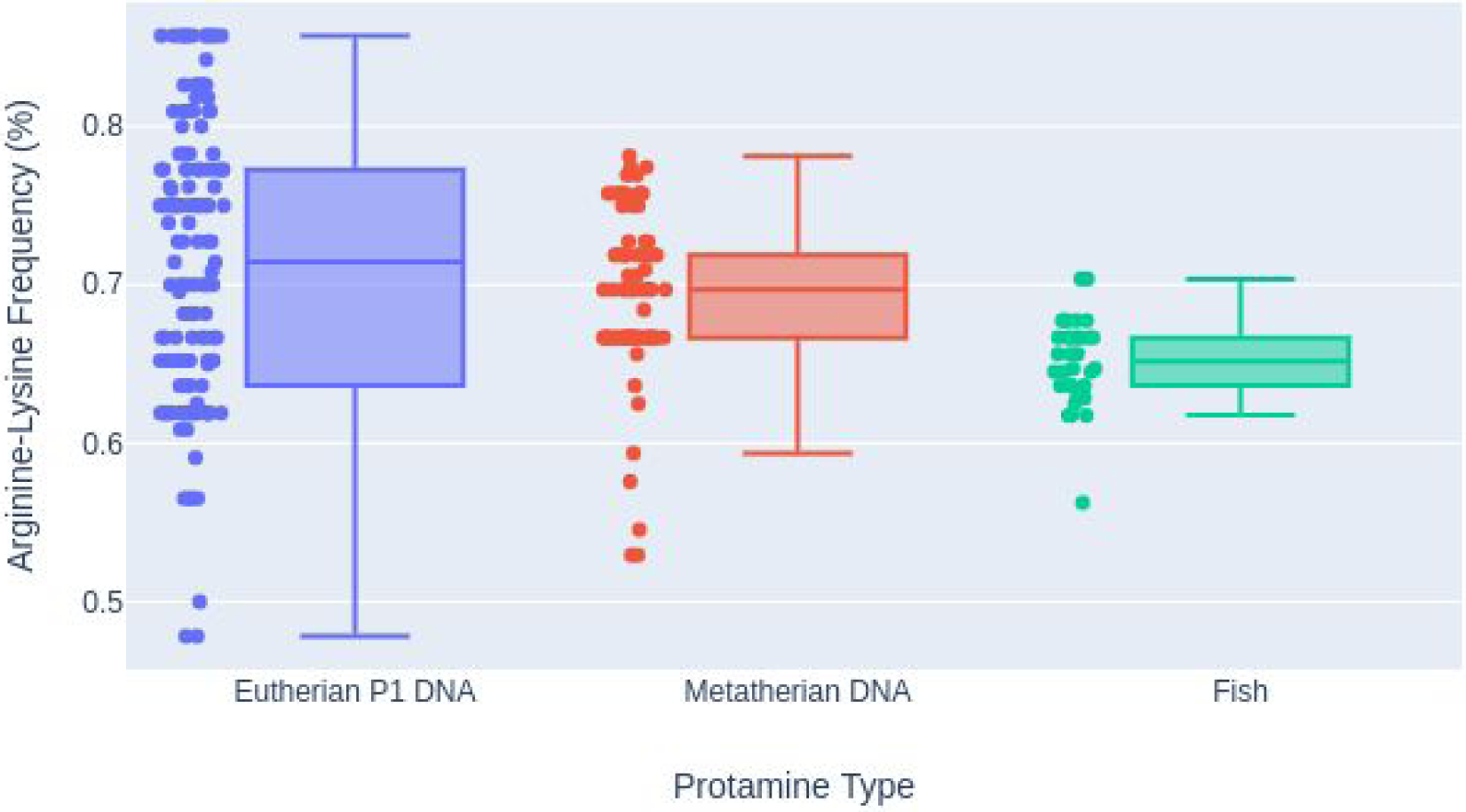
Box plots of the arginine-lysine frequency for the DNA binding regions of eutherian protamine P1, metatherian sperm protamine, and the whole sequences of fish protamines. Individual arginine-lysine frequencies are plotted next to the associated box plot.

## Discussion

The results from the MSA analysis of fish protamines showed no highly conserved residues above arginine. This likely indicates that the only functionally/structurally important residues in the fish protamines are arginine. The arginine-lysine density analysis showed that fish have a greater charge density across their entire protamines sequences than any of the other protamine groups.

The results from the MSA analysis of the eutherian protamine P1 sequences showed that the most highly conserved positions tend to be cysteine containing. The high evolutionary sequence conservation indicates that the positions are of great functional/structural importance. When these highly conserved positions are overlaid onto a proposed schematic structure for bull sperm protamine P1 [Balhorn et al. 1991; Vilfan et al. 2004], it is clear that the conserved positions align with the cysteines involved in intra- and intermolecular bonding in bull sperm protamine P1. It is also notable that the cysteines involved in the intramolecular cross-linkings were shown to be more highly conserved than those involved in the intermolecular cross-linkings. This likely supports the hypothesis that the hairpin-like secondary structure of eutherian sperm protamine P1s is required for proper DNA hypercondensation [Hutchison et al. 2017; Kirchhoff et al. 2019].

Comparing the metatherian P1 MSA to the eutherian P1 MSA, we find a number of commonalities. Both the N-terminal regions contain phosphorylation sites followed by blocks of arginine residues broken up by residues which can engage in cross-linking (cysteine in eutherians and tyrosine in metatherians) [Queralt et al. 1995]. A proposed schematic structure for metatherian sperm protamine P1 with conserved tyrosine positions is shown in Figure 7. The conserved tyrosine positions are visualized interacting in a similar cross-linking pattern as is observed in the cysteine containing eutherian mammal sperm protamines. Due to the similar conserved nature and similar spacing between the tyrosine residues in the metatherian protamine MSA and the cysteine residues in the eutherian protamine MSA, we hypothesize that metatherian protamines take on an analogous structure and folding mechanism as their eutherian counterparts. Folding of the metatherian protamines could possibly be facilitated by an orthologous enzyme to the glutathione peroxidase found in eutheria or an analogous peroxidase. There are specific peroxidases (e.g. certain myeloperoxidases) that are capable of catalysing dityrosine cross-linking in proteins [Bayse et al. 1972; Heinecke 2002; Mai et al. 2011]. However pi-pi stacking of two nearby tyrosines can represent another possible structural motif hypothesis [Lee et al. 2019].

**Figure 7.**
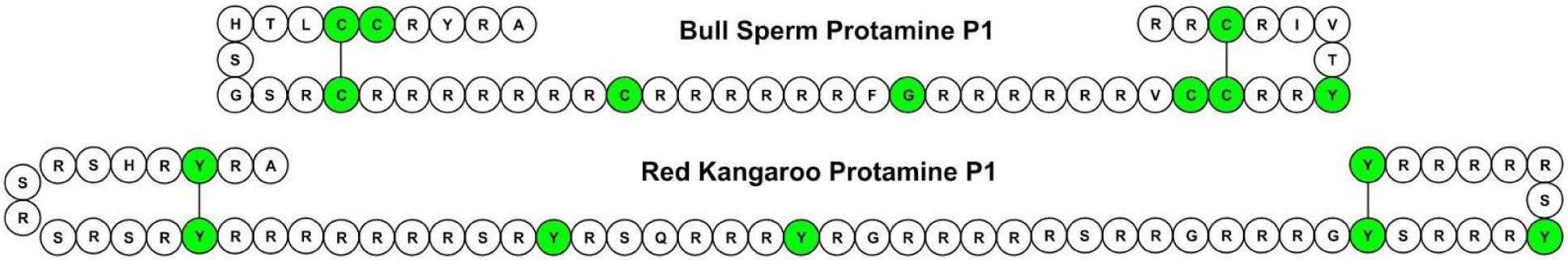
Highly conserved positions highlighted and overlaid onto proposed schematic structures for Bos taurus (Bull) and Macropus rufus (Red Kangaroo) sperm protamine P1s. Schematic structure of Bull P1 sperm protamines based off of Balhorn 1991 [Balhorn et al. 1991]. Red Kangaroo schematic structure assumed from similarity.

**Figure 8.**
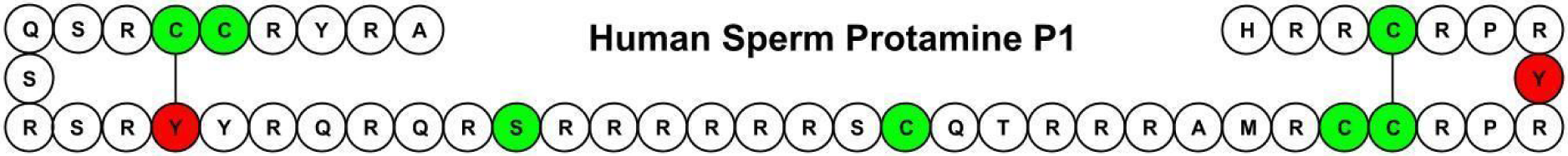
Highly conserved positions highlighted and overlaid onto proposed schematic structures for Homo sapien sperm protamine P1. Tyrosine containing positions highlighted in red and cysteine and other residue containing positions highlighted in green. Human schematic structure assumed from similarity [Balhorn et al. 1991].

The comparison of arginine-lysine frequencies of the differing protamine groups showed that although metatherian sperm protamines and fish protamines have similarly high charged residue frequencies across their entire sequences, metatherian mammals have an increased density of charged residues towards the middle of the sequences. This trend is also found in eutherian sperm protamine P1s where there is a central DNA binding region. The charged residue frequencies of the hypothesized DNA binding regions of eutherian P1 and metatherian sperm protamines were the most similar distributions by a significant factor.

Additional deviations are also apparent in the MSAs. One example is the tyrosine substitutions (tyr15cys and tyr16cys) of two positions in human sperm protamine P1, which are expected to be involved in intramolecular dicysteine cross-linking in the N-terminal staple fold. It is possible that cysteine-tyrosine cross-linking preserves the cross-linking function in human sperm protamines with these substitutions. Cysteine-tyrosine cross-linking, while not novel, is extremely rare in nature and are known to be mediated by copper metalloprotein enzymes [Martinie et al. 2012]. Also, prior research has shown that human spermatozoa are more susceptible to severe oxidative stress (>= 5 mM H_2_O_2_) in comparison to other eutherian mammals [Bennetts & Aitken 2005]. These substitutions could be an explanation for these observations.

Likewise, a counter deviation found in the metatherian alignment is with species of the *Planigale* genus, for which all species but *P. maculata maculata* contain cysteine substitutions in a number of highly conserved positions where tyrosine residues are expected (Supplemental Figure 5). It is notable that the change from tyrosine to cysteine is a single nucleotide substitution and that the arginine-lysine heavy regions are still found in the sequences of the *Planigale* species. What impact that these reciprocal substitutions may have on the fertility of the *Planigale* species and humans is not currently known.

## Conclusions

In summary, the common patterns of sequence conservation between eutherian and metatherian protamine P1 sequence families support hypotheses for dityrosine cross-linking in the metatherian P1 protamines and a rare cysteine-tyrosine cross-linking in human sperm protamine P1. The presence of cysteine cross-linking in a number of species of the *Planigale* genus also indicates that metatherian sperm protamines are likely capable of taking on a hairpin-like structure analogous to eutherian sperma protamine P1. This was additionally supported by the finding that metatherian sperm protamines also contain an increased density of arginine and lysine residues in the center of the sequence, which likely represents a large DNA binding region analogous to the DNA region found in eutherian sperm protamine P1 sequences. In addition to directly testing these hypotheses with wet lab and analytical experiments, the next logical steps involve searching for an analogous peroxidase enzyme with expression localized to the testis in metatherian mammals to provide further evidence of dityrosine cross-linking and possibly an analogous peroxidase with a copper binding cofactor or new mechanism to support the formation of a cysteine-tyrosine cross-linking in human sperm protamines. Moreover, these proposed mechanisms and structures may play a role in fertility, particularly human fertility.

## Methods

We used an entropy-based method to determine the functionally important residues in the MSAs of various protamine groups. Additionally, the charged residues densities of the protamine sequences in each group were determined. By comparing the conserved residues and the charged residue densities, we made predictions about structural features for related protamine groups and mechanisms behind their ability to bind DNA.

### Creation of Homologous Protamine Groups

All entries containing the keyword “protamine” were downloaded from the May 2019 release (release 2019_05) of the UniProt KnowledgeBase (SwissProt/TrEMBL) [UniProt 2017] to create the initial dataset of protamine and protamine-like proteins.

The protamine dataset was then broken down into four homologous groups based on existing UniProt gene name and organism classification annotations. The four groups were eutherian sperm protamine P1, eutherian sperm protamine P2, metatherian sperm protamine P1, and fish protamine. The eutherian sperm protamine P1 group was parsed by collecting all sequences which contained ‘Eutheria’ in their organism classification and the gene name of either ‘PRM1’ or ‘Prm1’. The same approach was performed for the eutherian sperm protamine P2 group, but using the gene name of either ‘PRM2’ or ‘Prm2’. The metatherian sperm protamine P1 group was parsed by collecting all sequences which contained ‘Metatheria’ in their organism classification and with the gene name of either ‘PRM1’ or ‘Prm1’. The fish protamine group was parsed by collecting any sequence which contained ‘Actinopterygii’ in their organism classification and that did not contain the word ‘like’ in their description. For the eutherian and metatherian groups only a single entry was allowed per organism per group. If multiple sequences existed in a single group, preference was given to the Swiss-Prot entry since these are reviewed entries. Therefore, each group is composed of orthologous genes, with the exception of the fish protamine group where some species have more than one protamine gene in the group.

### Additional Filtering and Truncation of Protamine P2 Sequences

Sperm protamine P2’s contain multiple post-translational cleavage sites, which lead to the removal of 40% of the amino terminus of these proteins [Balhorn 2007]. After processing, the protein sequence is slightly longer than that of protamine P1 and the processed protein of P2 also has a higher arginine frequency than that of the unprocessed sequence [Balhorn 2007]. As only the processed version of the protein interacts with DNA, the P2 alignment is truncated to only include the processed versions of these proteins. This is achieved by determining the closest post-translational processing site to each protein’s DNA binding region in the eutherian sperm protamine P2 alignment. The post-translating sites were found by using mouse sperm protamine P2 (MOUSE_PRM2) as a reference [Balhorn 2007]. MOUSE_PRM2 residue 44 is the closest post-translational processing site to the protein’s DNA binding region in mice. MOUSE_PRM2 residue 44 can be found at position 48 in the eutherian P2 sperm protamine alignment. Position 48 of the alignment was therefore used as the truncation site.

Aberrant sperm protamine P2 sequences (Rat, Boar, Bovin) caused gapping in an initial MSA and were found to lack significant translational expression in prior literature [Bunick et al. 1990; Maier et al. 1990]. Also the sequence for Chinese hamster was severely truncated. Therefore these sequences were removed before final alignment.

### Multiple Sequence Alignment and Conservation Analysis

A fasta file was generated for each homologous protamine group (i.e., eutherian P1, eutherian P2, and metatherian P1) and then MUSCLE 3.8.31 [Edgar 2004] was used to create an MSA using default settings.

Relative entropy (Kullback-Leibler divergence) was used to determine residue conservation scores for each position (column of residues) in the alignment. Relative entropy incorporates background frequencies of amino acids to measure the distance between the amino acid frequency in a position of the alignment versus the background frequencies [Capra & Singh 2007; Cover & Thomas 2006; Hannenhalli & Russell 2000].

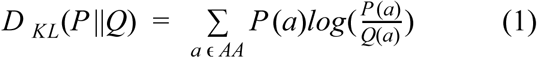

D(P||Q) is calculated for each position in the alignment and uses all 20 of the standard amino acids plus Asx (B) for Asp or Asn, Glx (Z) for Glu or Gln, and Xaa (X) for unknown. P(a) is the frequency of the amino acid in the position. Q(a) is the background frequency of an amino acid. For this analysis, the natural abundance of amino acids determined by the UniProt knowledgebase was used [UniProt Consortium 2017]. Relative entropy has been shown to be one of the most effective algorithms for determining functionally/structurally important residues from alignments and tied for the most effective method for determining positions playing a role in protein-protein interactions [Capra & Singh 1991; Hannenhalli & Russell 2000].

Additionally, a weighting was used to deal with the presence of gaps in the alignment. The gap weighting is incorporated by multiplying the calculated relative entropy measure by the percent of non-gap residues in the position.

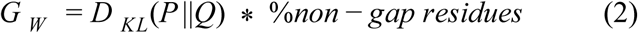

To determine which positions are conserved in the alignment, a conservation score threshold equal to a position entirely composed of arginine residues (∼4.1354) was used. If the gap weighted conservation score was greater than the threshold, the position was determined to be conserved. Methionine residues at the beginning of protein sequences are ignored.

### Arginine-Lysine Density Analysis

The arginine-lysine density of each protamine group was calculated by counting the number of arginine and lysine residues in each protamine sequence in the group. Histidine was left out of the analysis as it is mostly deprotonated at physiological pH. Lysine was included above a simple arginine frequency due to known DNA interactions for lysine residues in a variety of DNA binding proteins, but more importantly, an observed reduction in the severity of lower bound outliers for all protamine groups analyzed. This improvement in the lower bound outliers is greater for lysine than for the inclusion of any other amino acid (see Supplemental Tables 5-9). The quartile ranges for each group was then calculated and graphed using Plotly [Plotly 2015]. For eutherian sperm protamine P2, the truncated sequence was used, as determined by the method mentioned above. Additionally, for each protein in the eutherian P1 and metatherian sperm protamine alignments the charged residue density within the hypothesized DNA binding region was calculated. The hypothesized DNA binding region for the eutherian P1 alignment begins at position 17 and ends at position 46. The hypothesized DNA binding region for the metatherian alignment begins at position 16 and ends at position 56.

## Authors’ contributions

CDP performed the analyses. DCK, JED, and HNBM were all involved with the study’s conception and design. All authors read, edited, and approved the manuscript.

## Availability of Data and Material

All datasets, figures, and scripts are available through FigShare: https://doi.org/10.6084/m9.figshare.10292573.v1

## Supplemental Tables

**Table 1.**
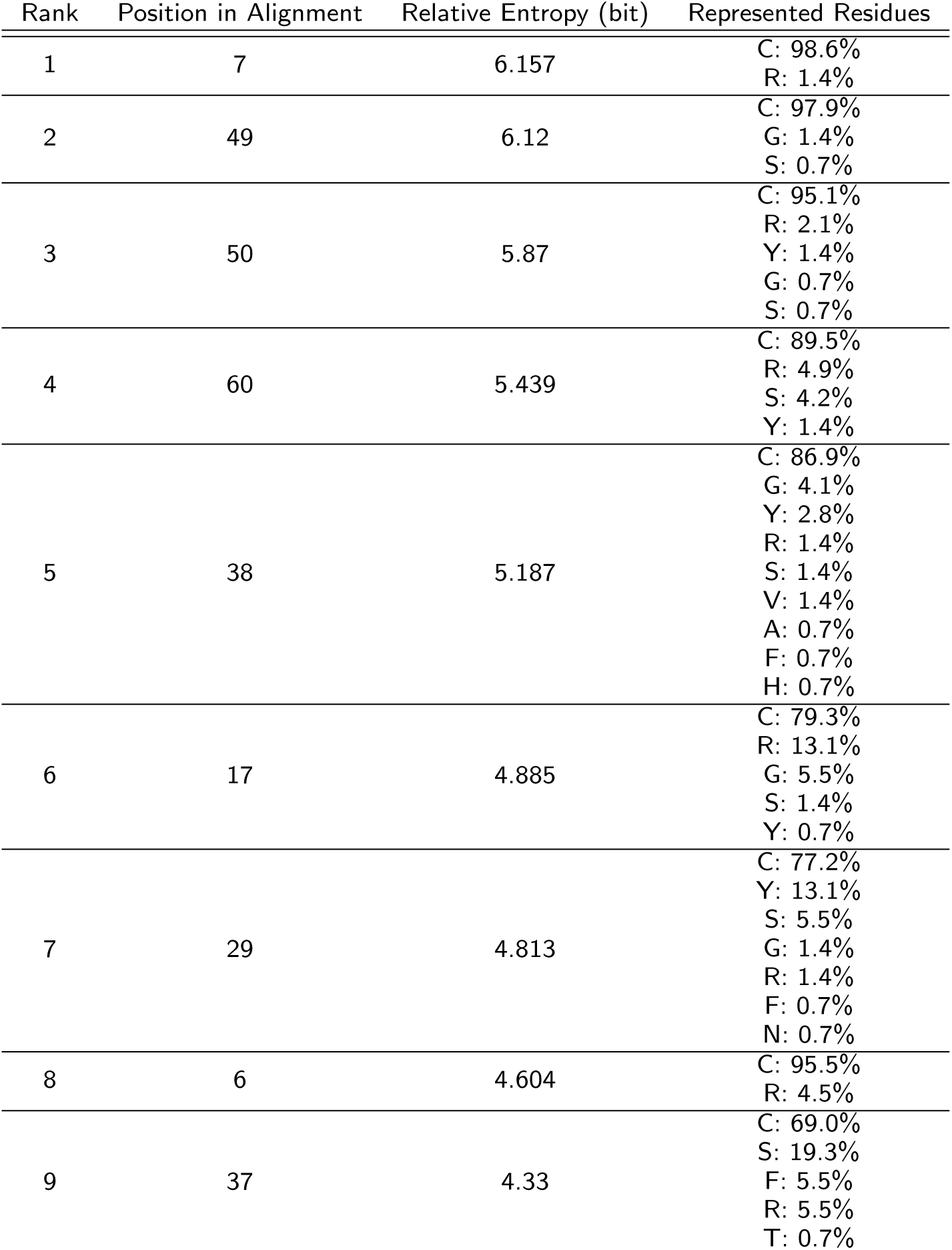
Highly Conserved Positions in Eutherian P1 Sperm Protamine MSA.

**Table 2.** Highly Conserved Positions in Truncated Eutherian P2 Sperm Protamine MSA.

| Rank | Position in Alignment | Relative Entropy (bit) | Represented Residues |
| --- | --- | --- | --- |
| 1 | 59 | 6.381 | C: 100.0% |
| 2 | 75 | 6.381 | C: 100.0% |
| 3 | 83 | 6.381 | C: 100.0% |
| 4 | 93 | 6.381 | C: 100.0% |
| 5 | 107 | 6.381 | C: 100.0% |
| 6 | 68 | 5.513 | H: 100.0% |
| 7 | 89 | 4.721 | H: 87.5%<br>N: 6.2%<br>P: 6.2% |
| 8 | 53 | 4.635 | H: 87.5%<br>G: 6.2%<br>S: 6.2% |
| 9 | 85 | 4.543 | H: 81.2%<br>Y: 12.5%<br>Q: 6.2% |
| 10 | 110 | 4.371 | H: 75.0%<br>Y: 18.8%<br>Q: 6.2% |
| 11 | 54 | 4.288 | Y: 56.2%<br>H: 43.8% |

**Table 3.**
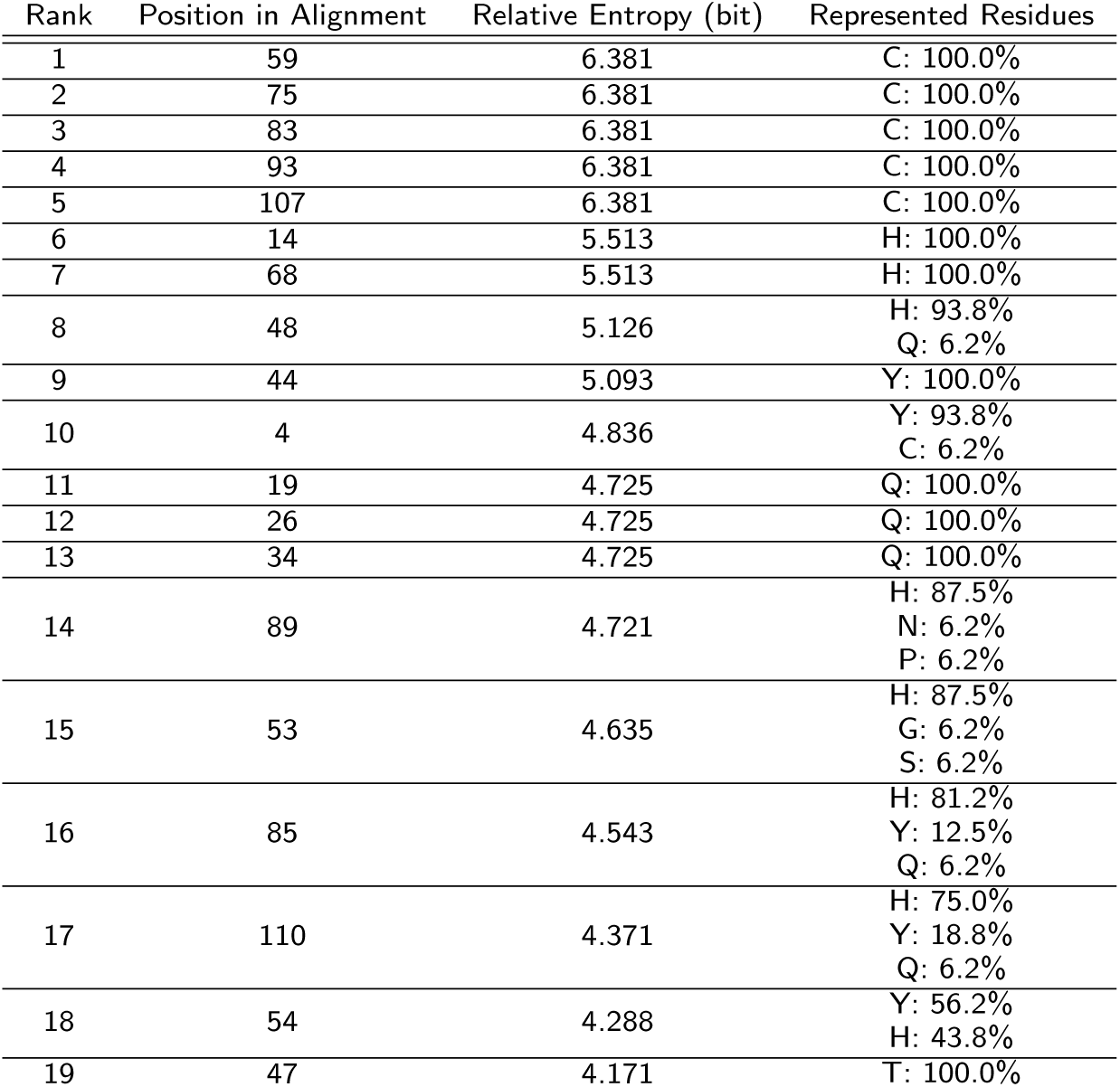
Highly Conserved Positions in Eutherian P2 Sperm Protamine MSA.

| Rank | Position in Alignment | Relative Entropy (bit) | Represented Residues |
| --- | --- | --- | --- |
| 1 | 59 | 6.381 | C: 100.0% |
| 2 | 75 | 6.381 | C: 100.0% |
| 3 | 83 | 6.381 | C: 100.0% |
| 4 | 93 | 6.381 | C: 100.0% |
| 5 | 107 | 6.381 | C: 100.0% |
| 6 | 14 | 5.513 | H: 100.0% |
| 7 | 68 | 5.513 | H: 100.0% |
| 8 | 48 | 5.126 | H: 93.8%<br>Q: 6.2% |
| 9 | 44 | 5.093 | Y: 100.0% |
| 10 | 4 | 4.836 | Y: 93.8%<br>C: 6.2% |
| 11 | 19 | 4.725 | Q: 100.0% |
| 12 | 26 | 4.725 | Q: 100.0% |
| 13 | 34 | 4.725 | Q: 100.0% |
| 14 | 89 | 4.721 | H: 87.5%<br>N: 6.2%<br>P: 6.2% |
| 15 | 53 | 4.635 | H: 87.5%<br>G: 6.2%<br>S: 6.2% |
| 16 | 85 | 4.543 | H: 81.2%<br>Y: 12.5%<br>Q: 6.2% |
| 17 | 110 | 4.371 | H: 75.0%<br>Y: 18.8%<br>Q: 6.2% |
| 18 | 54 | 4.288 | Y: 56.2%<br>H: 43.8% |
| 19 | 47 | 4.171 | T: 100.0% |

**Table 4.**
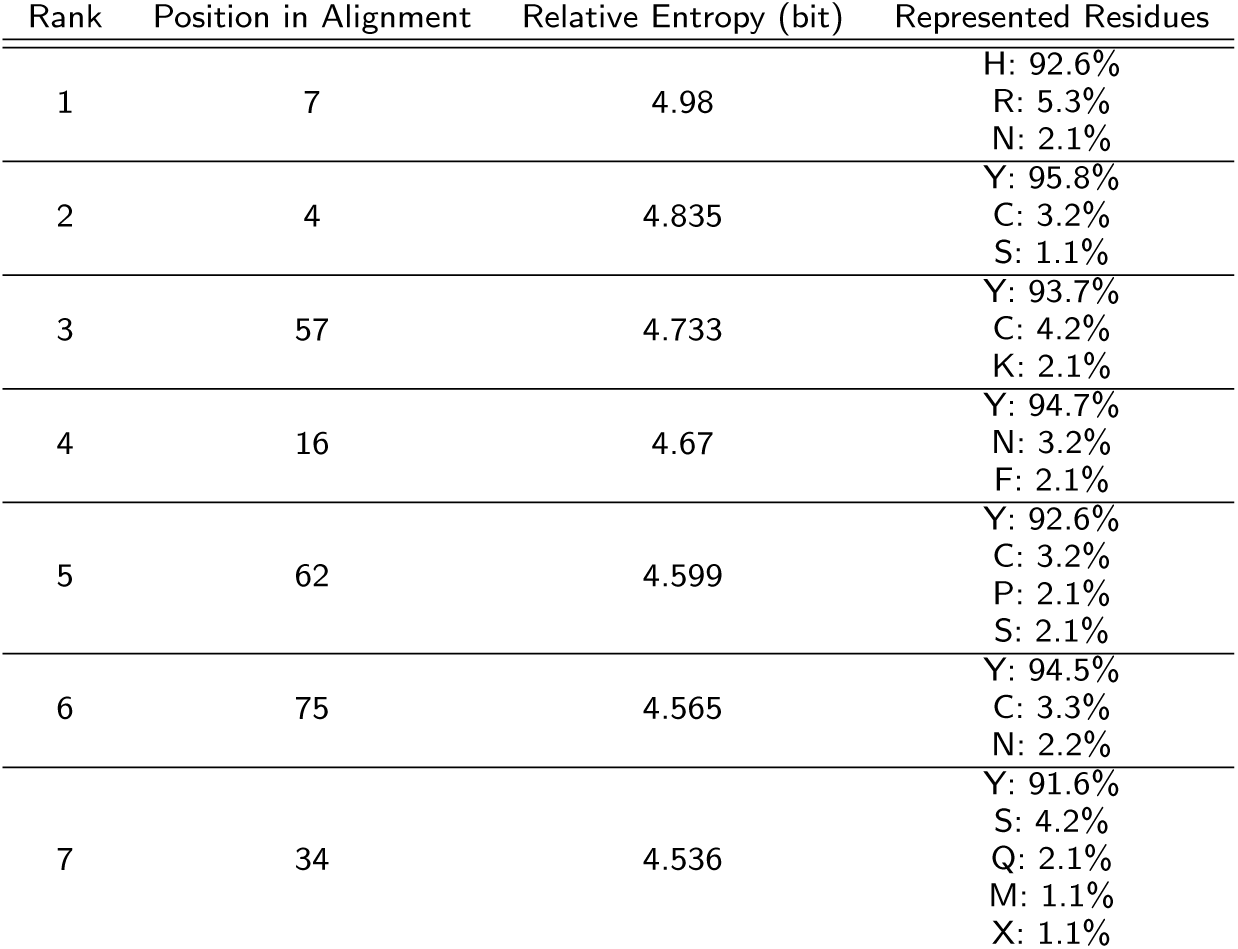
Highly Conserved Positions in Metatherian P1 Sperm Protamine MSA.

**Table 5.** Fish Protamines.

| Rank | Range | STD |
| --- | --- | --- |
| 1 | K 0.141 | K 0.027 |
| 2 | H 0.185 | H 0.035 |
| 3 | L 0.222 | L 0.039 |
| 4 | T 0.228 | T 0.041 |
| 5 | A 0.256 | E 0.045 |

**Table 6.** Eutherian P1 Sperm Protamines.

| Rank | Range | STD |
| --- | --- | --- |
| 1 | Q 0.182 | Q 0.034 |
| 2 | K 0.224 | P 0.044 |
| 3 | T 0.224 | T 0.046 |
| 4 | S 0.242 | N 0.046 |
| 5 | N 0.244 | A 0.049 |

**Table 7.** Eutherian P2 Sperm Protamines.

| Rank | Range | STD |
| --- | --- | --- |
| 1 | Q 0.07 | Q 0.017 |
| 2 | L 0.09 | T 0.022 |
| 3 | T 0.09 | L 0.023 |
| 4 | K 0.09 | E 0.024 |
| 5 | C 0.099 | V 0.025 |

**Table 8.** Truncated Eutherian P2 Sperm Protamines.

| Rank | Range | STD |
| --- | --- | --- |
| 1 | Q 0.116 | K 0.035 |
| 2 | K 0.117 | Q 0.037 |
| 3 | T 0.127 | T 0.04 |
| 4 | L 0.145 | L 0.044 |
| 5 | C 0.151 | G 0.045 |

**Table 9.**
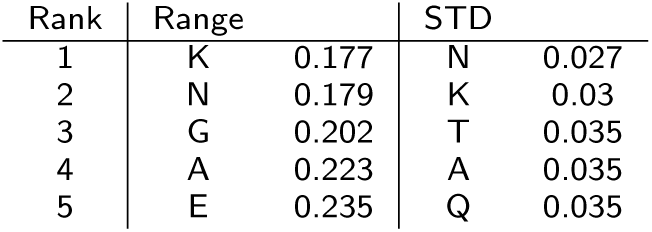
Metatherian Sperm Protamines.

**Supplemental Figure 1.**
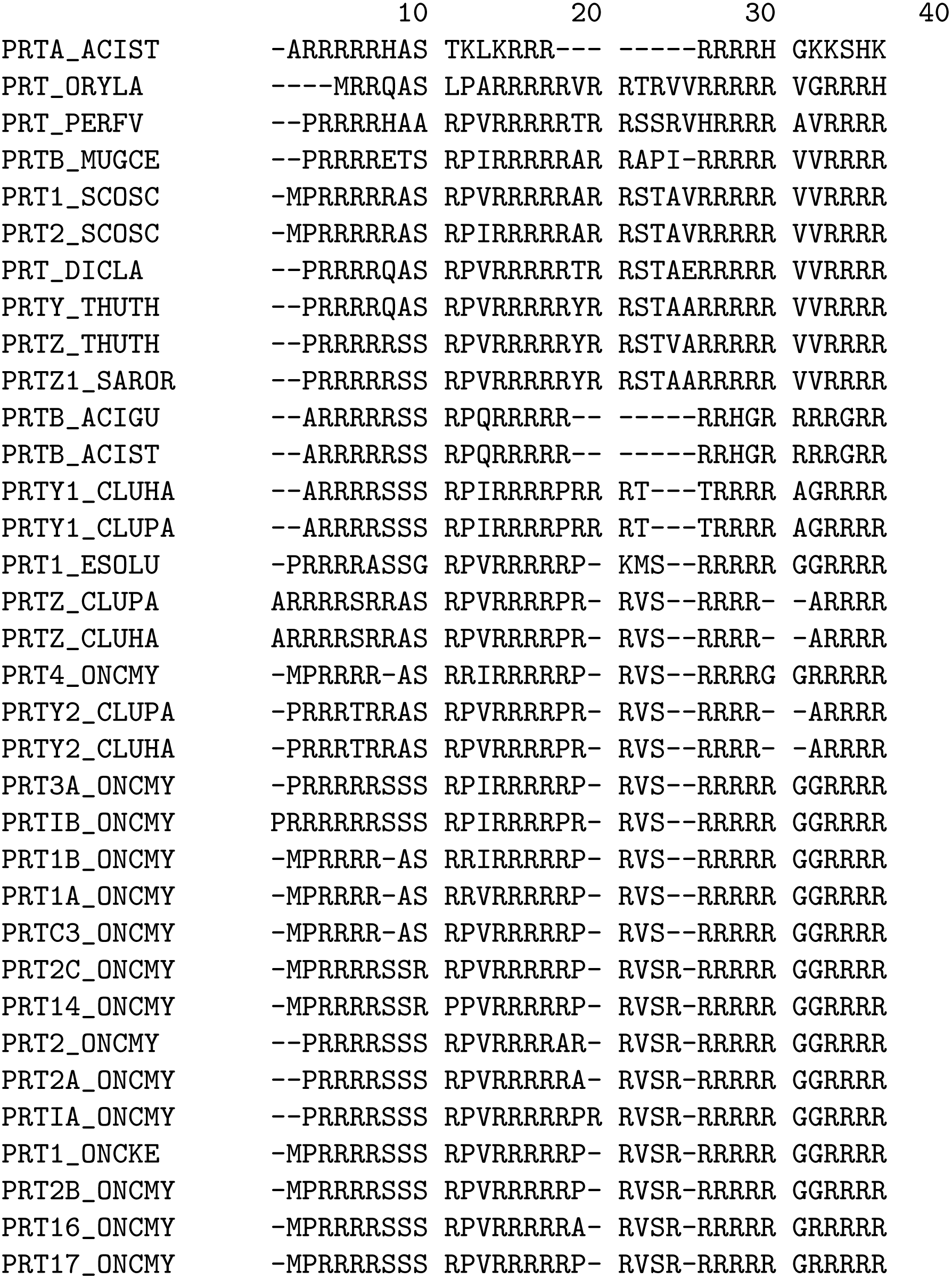
Alignment of fish protamines.

**Supplemental Figure 2.**
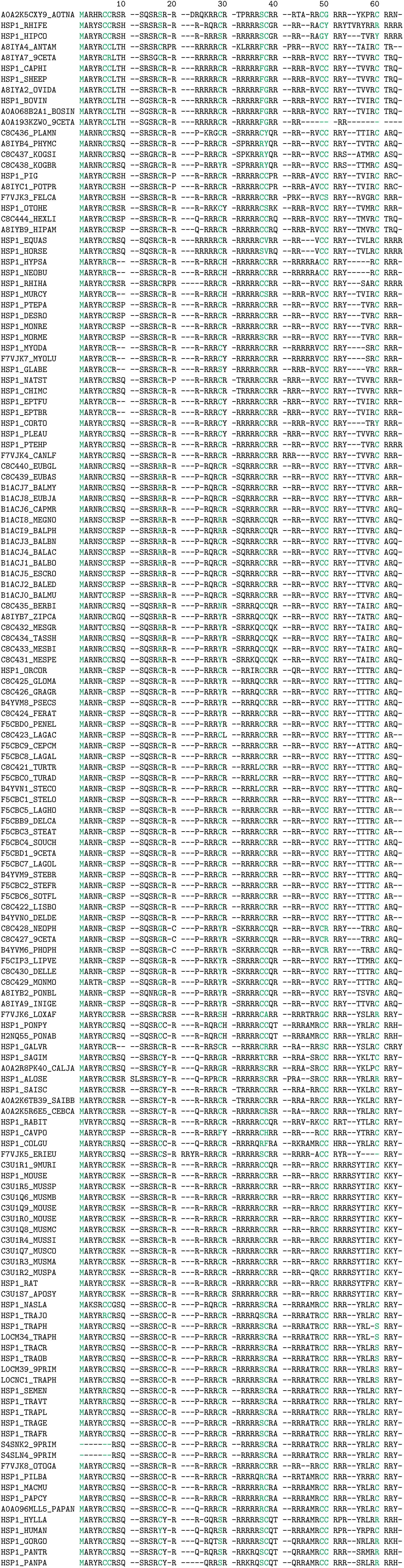
Alignment of Eutherian P1 type sperm protamines.

**Supplemental Figure 3.**
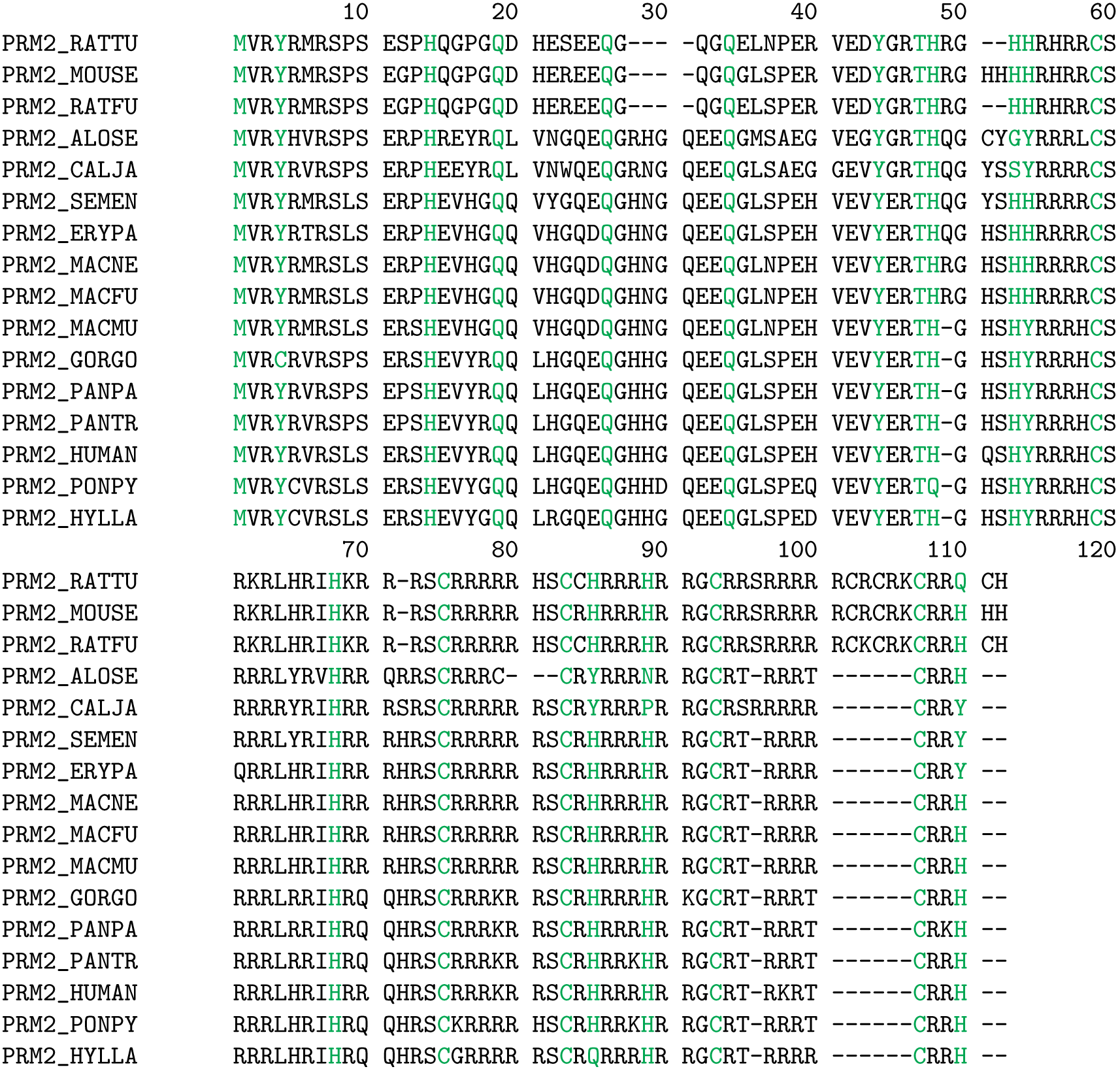
Alignment of Eutherian P2 type sperm protamines.

**Supplemental Figure 4.**
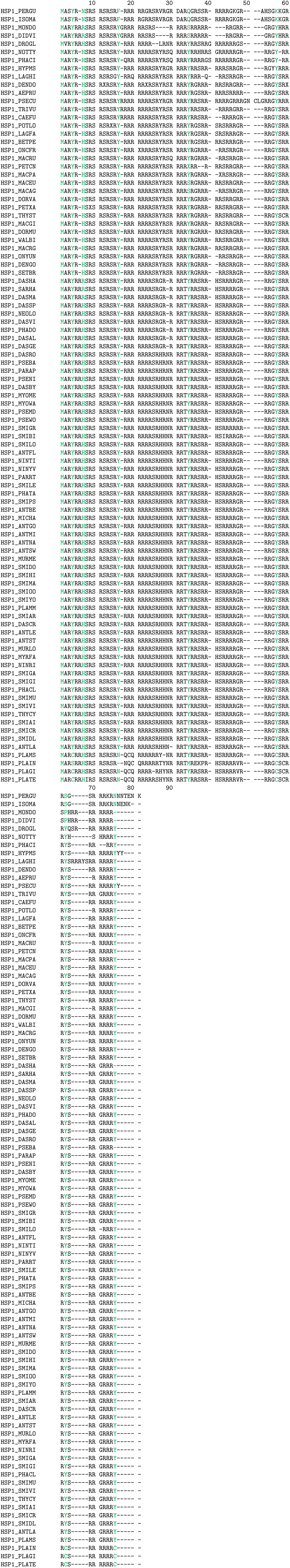
Alignment of Metatherian P1 type sperm protamines.

